# Improved yellow-green split fluorescent proteins for protein labeling and signal amplification

**DOI:** 10.1101/2020.09.27.315697

**Authors:** Shuqin Zhou, Siyu Feng, David Brown, Bo Huang

## Abstract

The flexibility and versatility of self-complementing split fluorescent proteins (FPs) have enabled a wide range of applications. In particular, the FP_1-10/11_ split system contains a small fragment that facilitates efficient generation of endogenous-tagged cell lines and animals as well as signal amplification using tandem FP_11_ tags. To improve the FP_1-10/11_ toolbox we previously developed, here we used a combination of directed evolution and rational design approaches, resulting in two mNeonGreen (mNG)-based split FPs (mNG3A_1-10/11_ and mNG3K_1-10/11_) and one mClover-based split FP (CloGFP_1-10/11_). mNG3A_1-10/11_ and mNG3K_1-10/11_ not only enhanced the complementation efficiency at low expression levels, but also allowed us to demonstrate signal amplification using tandem mNG2_11_ fragments in mammalian cells.

## Introduction

Fluorescent proteins (FPs), such as Green Fluorescent Protein (GFP), are a group of structurally homologous proteins that are widely used as genetically encoded fluorescent tags. Their structure usually consists of a cylindrical β-barrel, comprised of 11 β-strands, and a chromophore formed within the β-barrel. Splitting the β-barrel between strand 10 and 11 produces a spontaneously self-associating split fluorescent protein system, which we named as FP_1-10/11_ (1). Fluorescence depends upon complementation of FP_1-10_ and FP_11_ to form a mature FP. Starting from the green GFP_1-10/11_ (1), this toolbox has been expanded to different colors and fluorescent intensities by providing split versions of sfCherry (2), sfCherry2 (3), sfCherry3 (4) and mNeonGreen2 (mNG2) (3).

The self-complementing nature of FP_1-10/11_ systems has enabled a wide range of applications, including detection of protein expression (1), visualization of cell-cell interactions (3,5), and monitoring intracellular infection (6,7). In particular, the short fragment (16 amino acid) of FP_11_ can be easily integrated into any gene locus of interest with minimal disturbance of local genomic structure, enabling the generation of endogenously-tagged cell line libraries for the systematic analysis of protein function and localization landscape under almost-native conditions (8). Critically, tagging of low-abundance proteins presents a particular challenge because the fluorescent signal resulting from their weak expression is indistinguishable from background fluorescence. Unlike more abundant targets, for which live knock-in cells can be easily detected and sorted by flow cytometry, low-abundance targets usually require clonal expansion, duplication and genotyping by PCR, which is slower and more expensive. Using GFP_1-10/11_ labeling, approximately 30% of proteins expressed in HEK293T cells are above the detection threshold of flow cytometry cell sorting. The other 70% need stronger fluorescent signal to be distinguished from cellular auto-fluorescence (8). Under this scenario, improving the brightness of split-FP constructs is much desired to facilitate the generation of endogenously tagged cell line libraries, including low abundance proteins. Crucial enzymes and regulatory proteins like transcription factors are generally expressed at lower levels (9).

We have previously engineered a split mNG2_1-10/11_ system from the yellow-green fluorescent protein mNeonGreen (mNG), which is derived from *Branchiostoma lanceolatum.* The mNG fluorophore has a high extinction coefficient and quantum yield, making it 2.5 times as bright as EGFP (10). Accordingly, split mNG2_1-10/11_ showed significant brightness improvement compared to GFP_1-10/11_ for endogenous protein tagging. However, it remains dimmer than full-length mNG2 (3). We also note that high mNG2_1-10/11_ complementation signal relies on high expression of mNG2_1-10_ which may limit its application in sensitive cells.

The cellular brightness of a split FP_1-10_/11 system is shaped by two attributes: 1) the molecular brightness of the complemented protein and 2) the efficiency of complementation process. Analyzing split GFP_1-10/11_, mNG2_1-10/11_ and sfCherry2_1-10/11_, we have found that complemented split proteins have no significant difference in single-molecule brightness compared with their full-length counterparts (4). The reduced overall fluorescence signal comes from incomplete complementation of FP_1-10_ and FP_11_. Unlike GFP_1-10/11_, which has near perfect complementation behavior, mNG2_1-10/11_ and sfCherry2_1-10/11_ have relatively low affinity between the two fragments, leading to a sub-optimal complementation efficiency.

We previously developed sfCherry3_1-10/11_ by improving sfCherry2_1-10/11_ complementation efficiency. Here, we sought to engineer a brighter yellow-green colored FP_1-10_/11 with our understanding of complementation process and molecular brightness. One strategy was to improve the complementation efficiency of mNG2_1-10/11_ through directed evolution; Another was to leverage the outstanding complementation efficiency of GFP_1-10/11_ and boost its molecular brightness. Using these strategies, we have generated three new split FPs, mNG3A_1-10/11_, mNG3K_1-10/11_ and CloGFP_1-10/11_. The two mNG3 variants demonstrated increased fluorescent signal in *E. coli* and improved complementation at low-expression levels compared with mNG2_1-10/11_. CloGFP_1-10/11_ is no brighter than GFP_1-10/11_ despite saturating our screening platform. Finally, we demonstrate successful mNG3A/K signal amplification by tagging a target protein with five copies of the mNG2_11_ fragment in tandem. This approach can further boost overall fluorescence signal allowing in applications with low-expression targets or high background samples.

## Results

### Improving the complementation of split mNG2 using directed evolution

To improve the complementation efficiency of mNG2_1-10/11_, we performed directed evolution in an *E. coli* system using a strategy similar to our previous sfCherry3 work (4) and with more efficient cloning. In brief, mNG2_1-10_ and mNG2_11_ were expressed from a single mRNA with separate ribosome binding sites (RBS) (Fig 1A), so that the two fragments have a relatively constant expression ratio across all *E. coli* colonies. Besides the promoter to drive transcription, the RBS sequence also plays a critical rule in protein expression, and its efficiency can vary with the downstream genetic context. Here high protein expression was desired to maximize brightness differences between mutants.

**Fig. 1.**
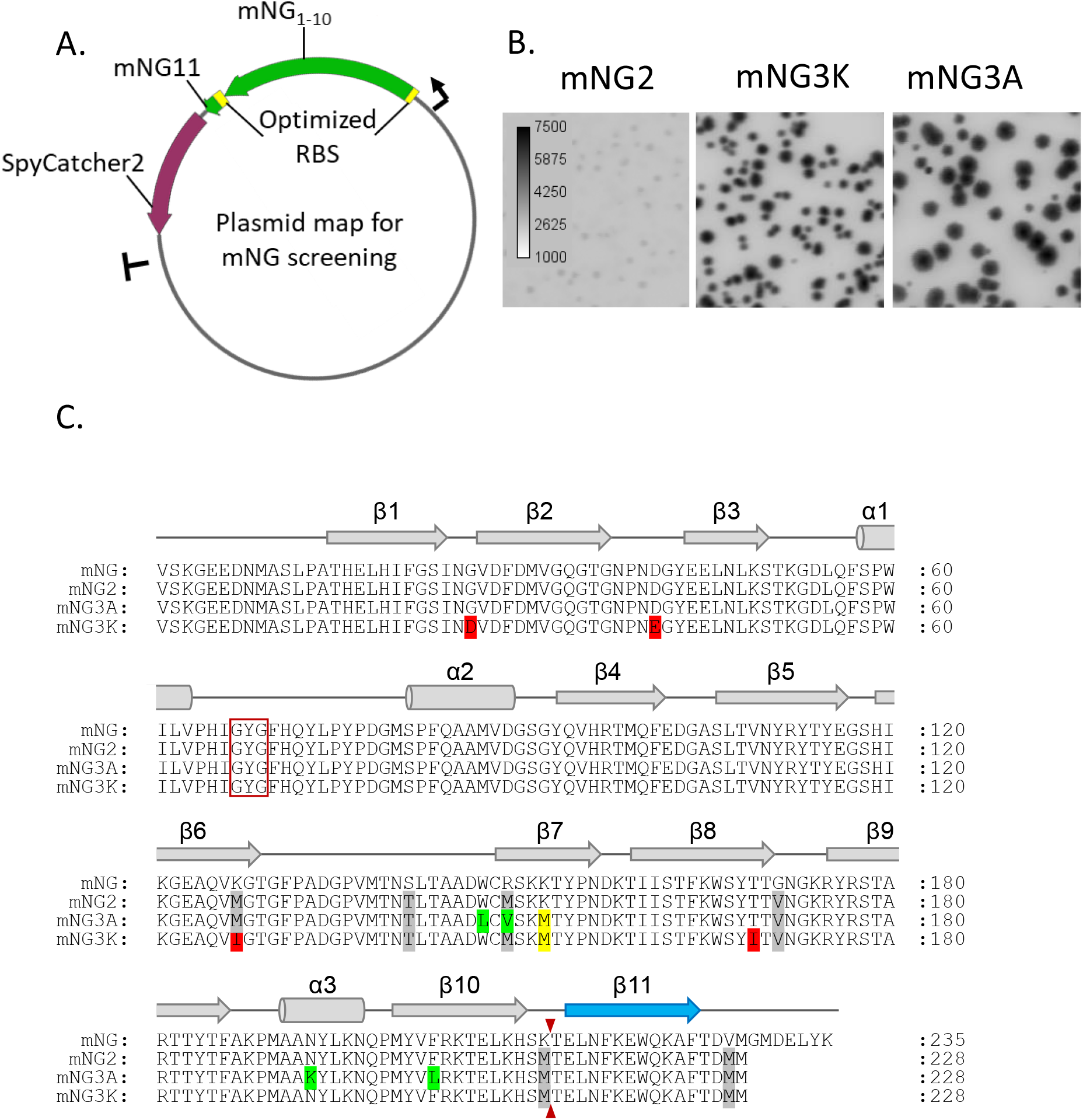
Identifying mNG variants with improved brightness in E. coli through directed evolution. A. Schematics of mNG screening platform; B. Fluorescence images of E. coli colonies expressing split mNG2, mNG3A and mNG3K; C. Protein sequence alignment of full length mNG, mNG2, mNG3A and mNG3K. Chromophore amino acids are indicated by a red box and the split site is indicated by the small red arrow. The position of α-helices and β-sheets are shown and numbered above the alignment.

We optimized the first RBS and second RBS (RBS2) for mNG2_1-10/11_ expression by screening a random 22 nt RBS library. Because small fragment peptides are prone to degradation, mNG2_11_ was fused to SpyCatcher2 (11) to improve its stability inside the cell. We used error prone PCR to introduce random mutations in mNG2_1-1-10_:RBS2:mNG2_11_ and this amplicon was integrated to our screening plasmid through golden gate assembly (Fig 1A). After four rounds of directed evolution and one round of DNA shuffling, the brightness of *E. coli* colonies grown on Luria broth (LB) plates increased substantially (Fig 1B).

Two candidates were identified, each with 5 substitutions in the FP_1-10_ fragment: W148L, M150V, K153M, N194K, F204L for mNG3A and G27D, D42E, M128I, K153M, T170I for mNG3K respectively (Fig 1C). Interestingly, the mNG3A mutations are clustered on the β-barrel, centering on strand 7 (Fig S2). Conversely, the mNG3K mutations are distributed around the β-barrel and are located towards the ends of β-strands or on the connective loops between them. Strand 7, which contains the K153M mutation common to mNG3A and mNG3K, is an irregular β-strand that hydrogen bonds to strand-10, potentially shaping its interface with the separate FP_11_ strand. Importantly, K153M represents a reversion to the wild type M143 at the equivalent residue in the parent protein LanYFP (10). The side chain of this residue projects into the β-barrel towards the fluorophore.

While characterizing these candidates more rigorously, we noticed point mutations in RBS2 which may have altered the expression level of mNG_11_ fragment. However, any contribution from the bacterial RBS used would be eliminated upon cloning into our mammalian cell expression plasmids.

### Engineering of CloGFP using rational design and directed evolution

Exploring an alternative approach to the production of a superior split FP, we sought to boost the molecular brightness of a split FP with efficient complementation. Split GFP has almost full complementation efficiency across a wide range of expression levels, so we want to take advantage of the high affinity of GFP_1-10/11_ system and improve its molecular brightness. Among the plethora of yellow-green fluorescent proteins developed from GFP, mClover, which was engineered from GFP to boost quantum yield, promised almost double the fluorescent brightness (12). Moreover, its photostability was further improved by 60% with N149Y, G160C, A206K mutations, in a variant named mClover3 (13).

Initially, we tested the effect of selected mClover mutations (L64F, T65G, Q69A, R80Q), which are close to the fluorophore, when introduced into a split GFP_1-10_ construct. However, the resulting sequence ‘CloGFP0.1’ produced essentially no fluorescence when cotransfected with GFP_11_::CLTA::P2A::mTagBFP in mammalian cells. We also tested the effect of introducing mClover3 mutations (L64F, T65G, Q69A, R80Q, N149Y, G160C, T203H, V206K), which are more broadly distributed through the β-barrel structure. Although this variant ‘CloGFP0.2’ lost more than 90% of the fluorescence signal compared to sfGFP_1-10_, some fluorescence was retained (Fig S1). This allowed us to use directed evolution to improve the cellular fluorescence signal.

Similar to the case of mNG3, we expressed the two fragments in *E. coli* using same screening plasmid under the same promoter (Fig 2A). We selected around 10 brighter mutants and used as a mixture of templates for next round of mutagenesis. Beneficial mutations accumulated in rounds of directed evolution, and most improvements appeared in the second and third rounds. In the 4th round no further improvement was observed, suggesting the screening system had reached saturation (Fig 2B). The brightest variant emerged in the 3rd round and contained G32Q, F46L, K101R, and S202L mutations, which we named ‘CloGFP’ (Fig 2C).

**Fig. 2.**
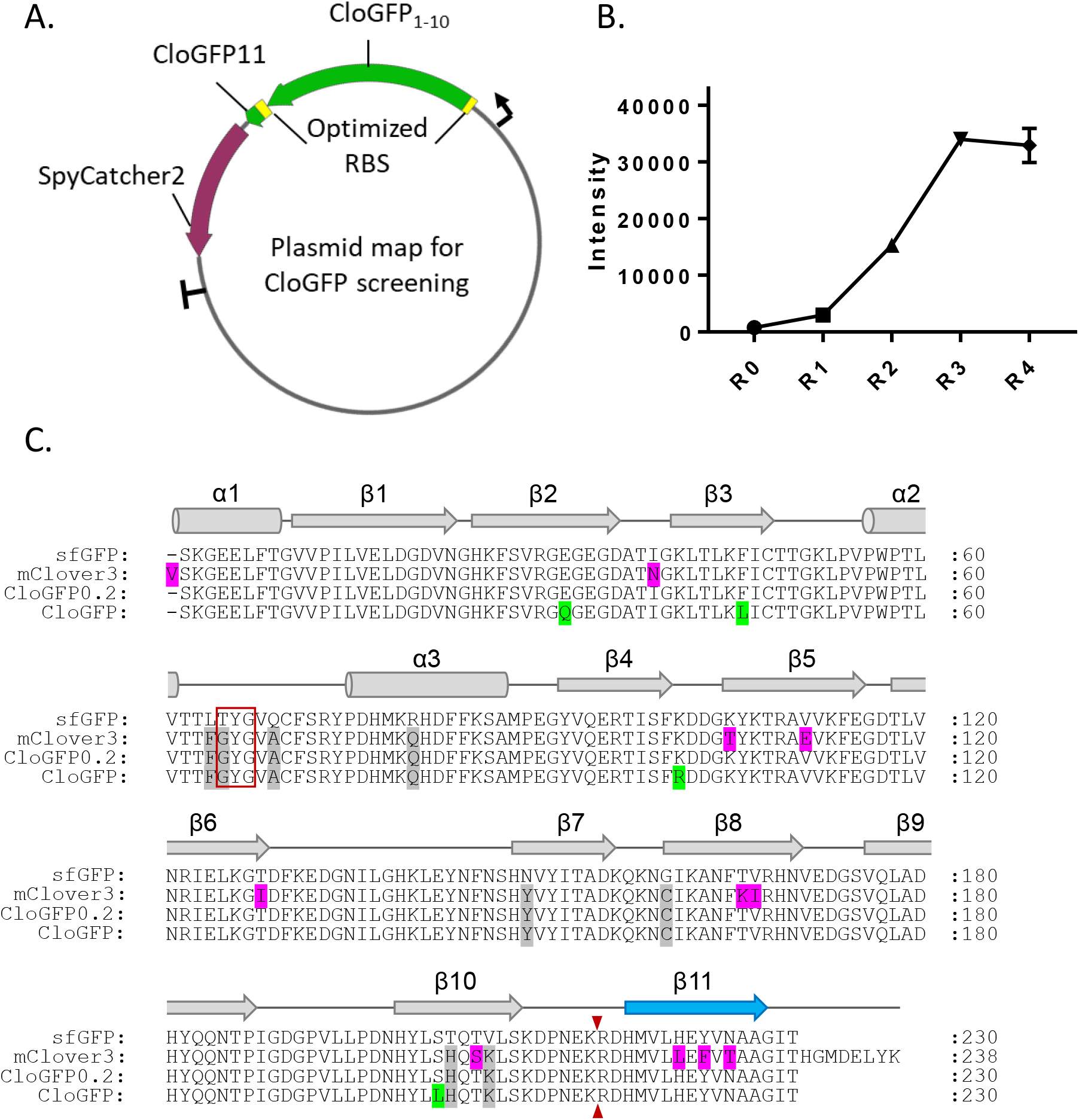
Identifying split CloGFP by recovery brightness through directed evolution. A. Schematics of split CloGFP screening platform; B. *E.coli* colony fluorescence intensity change during rounds of directed evolution, multiple colony with same mutant were measured in R4; C. Protein sequence alignment of sfGFP, mClover3, split Clover0 and split CloGFP. Chromophore amino acids are indicated by a red box and the split site is indicated by the small red arrow. The position of α-helices and β-sheets are shown and numbered above the alignment.

### Determining mammalian performance of mNG3 candidates and CloGFP

With mNG3(A/K)_1-10/11_ and CloGFP_1-10/11_ obtained by in *E. coli,* we investigated their complementation behavior in mammalian cells and whether their improved brightness is retained. We cloned mNG3(A/K)_1-10/11_ and CloGFP_1-10/11_ into a mammalian expression plasmid. mNG2_11_ and CloGFP_11_ were fused to the N-terminal of clathrin light chain A (CLTA), followed by TagBFP through a self-cleavable P2A site, so that the cellular concentration of mNG2_11_ and CloGFP_11_ could be quantified using TagBFP as the reference (Figs 3 and S3). We transiently transfected HEK293T cells with full length FP::CLTA::P2A::TagBFP or co-transfected with FP_1-10_ and FP_11_::CLTA::P2A::TagBFP, and analyzed the fluorescence from individual cells by flow cytometry (Fig 3).

**Fig. 3.**
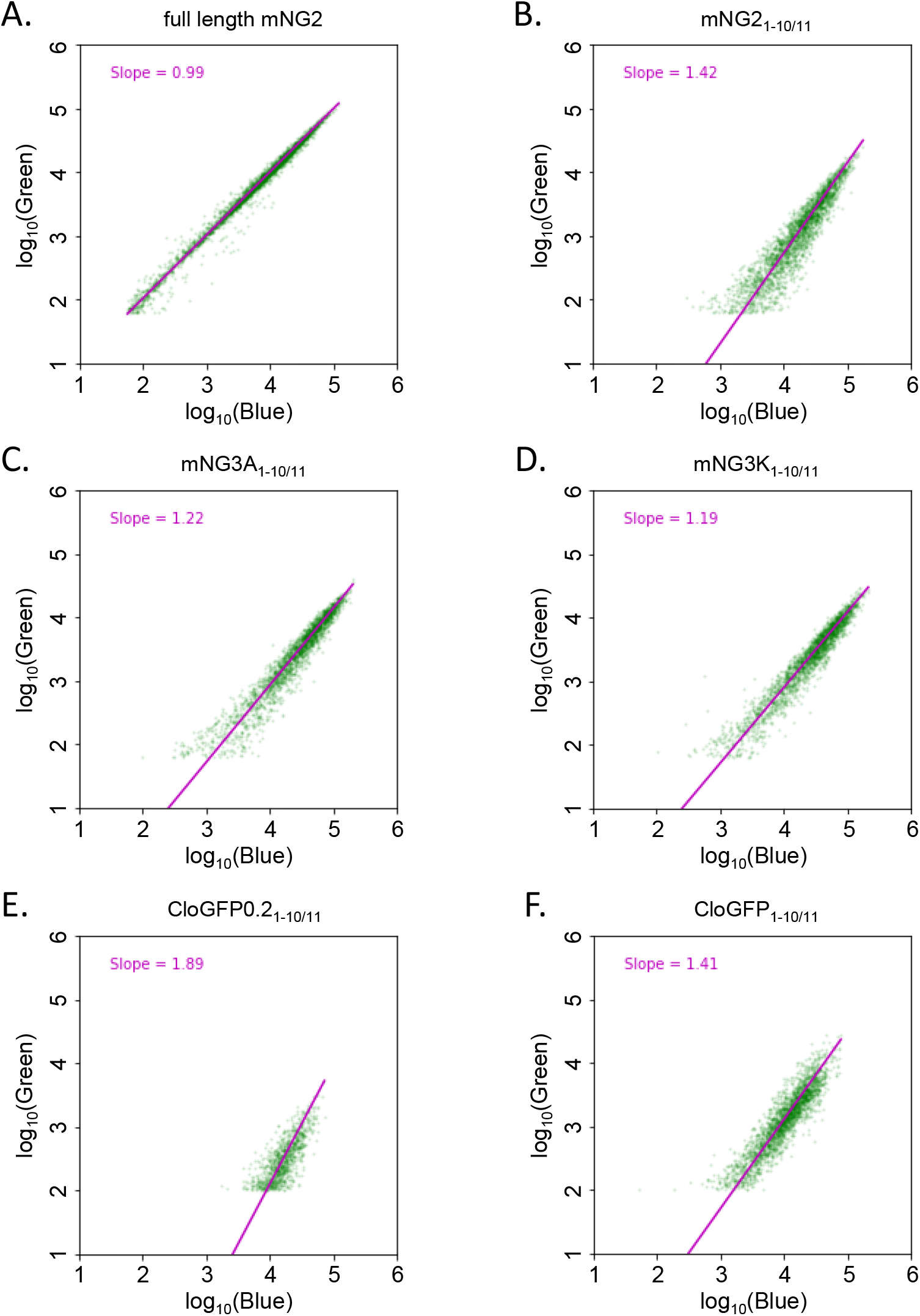
Improved affinity between FP_1-10_ and FP_11_ after engineering in mammalian cells. A-D, flow cytometry analysis of cell fluorescence by transient transfection in HEK293T expressing A. full length mNG2, B. mNG2_1-10/11_, C. mNG3A_1-10/11_, and D. mNG3K_1-10/11_. E-F flow cytometry analysis of cell fluorescence by transient transfection in HEK293T expressing E. CloGFP0_1-10/11_ and F. CloGFP_1-10/11_. The data are normalized to a full-length control.

The full-length mNG2::CLTA::P2A::TagBFP control yielded a linear relationship between mNG2 intensity in the Green channel (488 nm excitation) and TagBFP intensity in the Blue channel (405 nm excitation). In the log-log plot, a linear fit to this relationship gave a slope of almost exactly 1 (0.99) (Fig 3A), validating the use of P2A site to produce proportional concentrations of mNG2 and TagBFP across the range of expression levels resulted from transient transfection. For the split FP_1-10/11_ systems, on the other hand, it has been previously shown that the complementation process begins with a dynamic binding equilibrium between the two fragments (4,14). When the effective *K_D_* is much higher than the concentrations of co-transfected FP_1-10_ and FP_11_ fragments, this equilibrium cause the complemented FP signal to depend linearly on the expression reference with a slope of 2 on a log-log plot, while lowering the *K_D_* reduces this slope to 1 as the complementation becomes more efficient and approaches saturation (4). Therefore, we could use the slope from linear fit to characterize the effective *K_D_* relative to the concentration in our flow cytometry experiments. Compared to split mNG2_1-10/11_ which had a slope of 1.42 (Fig 3B), the new mNG3A_1-10/11_ and mNG3K_1-10/11_ had smaller slopes of 1.22 and 1.19, respectively (Figs 3C-D), indicating that they have lower effective *K_D_*, which resulted more efficient complementation and higher complemented fluorescence signal at the tested expression levels. These results confirmed the performance improvements in mammalian systems for these constructs engineered in *E. coli*.

For the variants incorporating mClover3 mutations into split GFP_1-10/11_, we used sfGFP::CLTA::P2A::TagBFP as the control (Fig S3). The initial CloGFP0_1-10/11_ yielded a slope of 1.89 (Fig 3E). This suggests that the introduction of the mClover3 mutations into GFP_1-10/11_ severely compromised its otherwise highly efficient complementation (Fig S3). With improvements from directed evolution, CloGFP_1-10/11_ produced by directed evolution gave a slope of 1.41, comparable to that of the mNG2_1-10/11_. We compared the performance of CloGFP_1-10_ with or without a mutation in the FP_11_ fragment (Y9F) which arose during the directed evolution. Yet the presence of absence of 4-hydroxyl group at this position had no effect in our assay (Figs S3 I and L). Ultimately, CloGFP_1-10/11_ was outperformed by mNG3A/K_110/11_, with the latter having lower effective *K_D_* (higher complementation efficiency) as well as overall stronger complemented fluorescence signal.

### Signal amplification using tandem mNG_11_

Besides improving complementation or single molecule brightness, another strategy to boost the apparent fluorescent intensity is to tag proteins of interest with multiple fluorescent proteins linked in tandem. Using a split-FP system, this can be achieved without repetition of the full FP sequence which can be difficult to clone and verify by sequencing and can perturb expression or localization of the tagged protein. This approach was used for single particle tracking of an intraflagellar transport (IFT) protein in primary cilia with IFT20::GFP_11_×7 (2) Tandem sfCherry_11_ tags labeling β-tubulin or β-actin also demonstrated enhancement proportional to the number of tandem repeats (2).

In preliminary experiments (not shown), however, we observed that mNG2_1-10/11_ signal could not be amplified using a tandem FP_11_ strategy. Here we wanted to see if our new mNG3A/K splits can be applied to single amplification by tandem repeats. We generated plasmids expressing either mNG2_11_ or a 5x tandem array of mNG2_11_, fused to the histone H2B protein. Each mNG2_11_ array length was co-transfected into HeLa cells with one of our mNG2, mNG3A_1-10_ or mNG3K_1-10_ constructs, allowing us to compare their brightness in the cell nuclei (Fig 4). With mNG2_1-10_, 1x and 5x samples showed no significant difference. In contrast, mNG3A_1-10_ and mNG3K_1-10_ produces 1.5-fold and 2.3-fold improvements, respectively, in the median cell signal for 5x mNG2_11_. Although this is still not linear amplification at 5x, it is a substantial improvement compared to no amplification in the mNG2_1-10/11_ case.

**Fig. 4.**
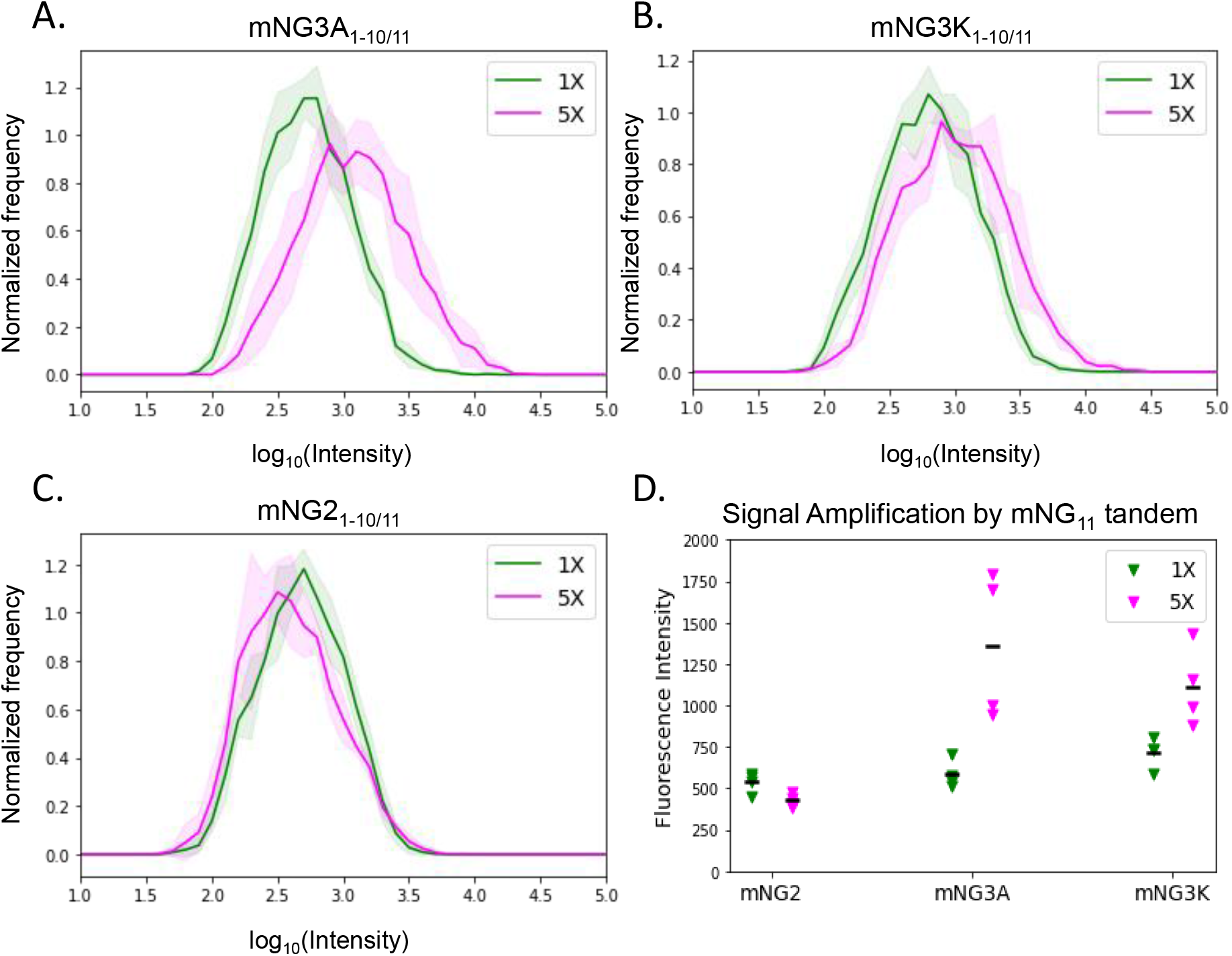
Significant signal amplification by tandem mNG_11_ with mNG3A_1-10_ and mNG3K_1-10_. A-C. Fluorescence intensity distribution of individual cells transiently cotransfected with FP_11_ (1x mNG_11_ in blue or 5x mNG_11_ in green) and FP_1-10_ (mNG2_1-10_, mNG3A_1-10_ or mNG3K_1-10_), each condition has four replicates, solid lines show the mean of four distributions, the shadow indicates standard deviation. Representative images of the cells are shown in Figs S4 and S5. D. Mean intensity comparison between the same conditions as A-C.

## Discussion

Here we used directed evolution combined with rational design to improve the brightness of two yellow-green split-FPs. This work revealed the specific merits of complementation-, or brightness-centered approaches to increase split-FP brightness, and yielded three new yellow-green split-FPs. In the context of split-FP complementation efficiency, our tandem tag experiments have important implications for endogenous tagging and detection of low expression proteins.

We chose *E. coli* as an initial screening system for their fast reproduction and high transformation efficiency. Plating *E. coli* in agar plates, we can screen up to 6000 mutants per round in two days. Although fluorescence intensity can be measured directly in this format, the readout is intrinsically linked to protein expression level, which depends on factors such as plasmid copy number and colony size. In our screen, both FP_1-10_ and FP_11_ fragments were encoded on a single plasmid to minimize variation in their relative expression levels. Two ribosome binding sequences (RBS) were used to drive translation of FP_1-10_ and FP_11_::SpyCatcher2 from the same mRNA. By removing the possibility of *E. coli* transformed with one fragment but not its complement, our screen is more sensitive to beneficial mutations which are less likely to be missed due to incomplete transformation. During mNG3 and CloGFP engineering, we found that mutations increasing the activity of RBS2 frequently arise in a single round of screening because they directly amplify FP_11_ expression. This makes it difficult to compare and quantify brightness across mutants without re-cloning them behind the original RBS. Emerging strategies for directed evolution in mammalian cells are potential solutions to this limitation(15,16). Spontaneous mutagenesis in continuous style may further boost its screening efficiency with high throughput and minimal human intervention(17).

As we and others continue to develop and improve methods to optimize the cellular brightness of split FPs, this study provides valuable insights as to which strategies work best. Our optimization of split mNG produced two new variants, sharing a common K153M reversion. It is likely that prior mutagenesis to maximize single molecule brightness of mNG did so at the expense of stability at the FP_1-10/11_ interface. In our work, this appears to have been reversed by restoration of the original methionine at residue 153, accompanied by a cluster of proximal compensatory mutations in mNG3A or a more distal set in mNG3K.

Interestingly, the degree of improvement achieved was similar for both mNG3A and mNG3K despite little overlap in the mutations. This may suggest that potential improvements to mNG2_1-10_ complementation exist as mutually exclusive sets of mutations, reflecting the tension between brightness and complementation efficiency characteristics. The near perfect complementation of split-sfGFP suggest that sfGFP is well suited to the 1-10/11 split site. However, for divergent 11-stand beta-barrel sequences, variant residues that improved single molecule brightness of the folded protein, frequently reduce the complementation efficiency. Further improved variants of split mNG system may require a priori determination of the optimal split site which may be combined with circular permutation to retain the benefits of a short peptide tag.

Unlike sfGFP_1-10_, mNG2_1-10_ has negligible non-complemented background, allowing it to be expressed at a high level to drive complementation efficiency without impacting the fluorescent signal to noise ratio (4). While we have not yet observed significant negative effects of overexpression on cell physiology, it remains a concern, especially for sensitive cells and applications. The improved affinity of our split mNG3 variants allow detection at lower levels of mNG_1-10_ expression, reducing potential perturbation to the cells. The enhanced ability of mNG3A_1-10_ or mNG3K_1-10_ fragments to occupy tandem FP_11_ repeats, also makes detection of tandem mNG2_11_ with our mNG3 variants a viable alternative to a tandem sfGFP_11_ system. Using mNG3 may be preferable when supplying high levels of mNG3_1-10_ is readily achievable, but intrinsic background fluorescence from sfGFP_1-10_ presents a problem. This is the situation when screening cell populations for successful knock-in of tandem FP_11_ repeats on a low abundance protein target by fluorescence activated cell sorting.

CloGFP_1-10/11_ has a lower complementation efficiency than sfGFP_1-10/11_, but produces a similar cellular brightness, consistent with a higher expected molecular brightness. For detection of proteins endogenously tagged with an FP_11_ by transient transfection CloGFP may outperform sfGFP, because the overexpression of CloGFP_1-10_ can drive the equilibrium towards 100% complementation, while each complemented FP will be brighter. CloGFP may also prove valuable for single molecule studies, where molecular brightness is more important than 100% labelling efficiency. Conveniently, CloGFP and sfGFP share the same FP_11_ sequence, allowing the FP_1-10_ fragment to be chosen as appropriate. For example, transfection with sfGFP_1-10_ might be used to study the macro structure of FP_11_-tubulin in microtubules, while a CloGFP_1-10_ transfection could be used to track individual FP_11_-tubulin molecules over time.

## Materials Methods

### Directed evolution

We used random mutagenesis and screening to improve the brightness of mNG2 and split CloGFP0.2 as previous described (3,4). The amino acid sequence and split of mNG2 was from our previously published paper (3). The mClover or mClover3 mutations (12,13) were merged into split sfGFP_1-10_ and synthesized directly from IDT. They were subjected to random mutagenic PCR using GeneMorph II Random Mutagenesis Kit (Agilent Technologies) to generate a library of candidates. PCR product were gel-purified and ligated into *E. coli* expression plasmid as in Fig1A or Fig2A using Golden Gate Assembly. The plasmid pools were mixed with Phage Display Electrocompetent TG1 Cells (Lucigen) and transformed by electroporation with the Gene Pulser Xcell™ Electroporation Systems (BioRad). Cells were recovered in SOC for 20 min at 37°C after electric pulse. Then cells were evenly plated on LB-agar plates with 50 μg ml^−1^ Kanamycin (Teknova) with glass beads for overnight incubation at 37°C. Proper density was needed to maximize screening capability and get well-isolated colonies at the same time. Plates with *E. coli* colonies were imaged at green channel using a BioSpectrum Imaging System (UVP) or Typhoon gel imager (GE Lifescience). Around 30 plates were imaged each round screening ~10,000 colonies. To further minimize brightness variation from plasmid copy number or colony size, streak plates were made from the brightest colony of each plate. After isolating and sequencing those colonies, the brightest ones with least RBS mutations were used as templates for next round of directed evolution.

### Mammalian Cell Culture and Transfection

Human HEK 293T cells were cultured in Dulbecco’s modified Eagle’s medium (DMEM) with high glucose (Gibco), supplemented with 10% (vol/vol) Fetal Bovine Serum and 100 μg ml^−1^ penicillin/streptomycin (Sigma-Aldrich) and maintained at 37°C and 5% CO2 in a humidified incubator. Cells were seeded at 4 × 10^4^ cells per well of a 48-well plate one night before transfection. For split proteins, 90 ng FP_11_ and 180 ng FP_1-10_ plasmids were mixed in OptiMEM and for full length, 180 ng plasmid, 0.8 μl FugeneHD (Promega) were used. Cells were harvested for flow cytometry 48 hours after transfection. For imaging, 8 well glass bottom chambered Lab-Tek (Nunc) was first coated with fibronectin (Sigma Aldrich) for 45 min, followed by three washes with PBS to help cells attach better. Similarly, cells were seeded at 5 × 10^4^ cells per well one night before transfection and 120 ng FP_11_, 240 ng FP_1-10_ plasmids, 1.08 μl FugeneHD reagent were used. After 48 hours, cells were fixed with 4% paraformaldehyde (diluted from 16% paraformaldehyde aqueous solution, Electron Microscopy Sciences) for 15 min and washed with PBS for three times. Then cells were imaged on a Nikon Ti-E inverted wide-field fluorescence microscope equipped with an LED light source (Excelitas X-Cite XLED1), a 10x air objective (Nikon PlanFluor 0.3 NA), a motorized stage (ASI), and an sCMOS camera (Hamamatsu ORCA Flash 4.0).

### Flow Cytometry

.fsc files containing the raw flow cytometry data were read into python using the [fscparser](https://github.com/eyurtsev/fcsparser) package. Detected events were filtered to remove cases that saturated one or more recorded channels (Fig S3A). Unsaturated events were then ‘Scatter’ gated based on the forward and side scatter channels (FSC-A and SSC-A respectively) to remove abnormal or dying cells and non-cellular debris (Fig S3B). Events within the Scatter gate were further screened to remove ‘doublet’ events, where a single droplet produced by the flow cytometer is likely to contain more than one cell (Fig S3C). The fluorescence measurements from filtered events were then transformed onto a log10 axis for analysis. Fluorescence thresholds were set based on a control sample of untransfected cells (Fig S3D). Gradients reflecting the split FP complementation efficiency were calculated as a linear regression, however as we have noted previously, certain preprocessing steps are necessary to fit the flow cytometry data appropriately (4). Fluorescence channel measurements each interrogate different fluorescent proteins and were performed at different wavelengths and with differing laser intensities. To account for these sources of variation, we fit a control sample expressing an unsplit version of our split-FP fused to the reporter FP (e.g. mNG-TagBFP). Using this fit, we rescaled our data such that a 1:1 relationship between complementation and the reporter FP yields a fixed gradient of 1. Variations in transfection efficiency can also affect the estimated gradient.

The python code used for this analysis was written to facilitate investigation of split FP brightness and complementation and is available on GitHub (https://github.com/BoHuangLab/altFACS).

### Image analysis

Images were stored as Tiff files and opened with ImageJ. To segment individual cells in the image, we calculated a binary mask. First, the background was subtracted from the green fluorescence image using the ImageJ Subtract Background function with rolling ball radius at 20 pixels. A Bandpass filter was applied to enhance cell edges to create cell masks. Particles of size 80-2000 μm^2^, circularity 0.1-1.0 were analyzed, and median value of each particle was used as intensity of each cell. The ImageJ Macro used for image processing are available on GitHub (https://github.com/BoHuangLab/ImageJ-macro-for-cellular-intensity-analysis). Cell intensity distribution was plotted as histogram in python. Each condition had four replicates and their histograms were averaged. The python script for these plots can also be found on GitHub (https://github.com/BoHuangLab/ImageJ-macro-for-cellular-intensity-analysis).

## Acknowledgements

Thank you to our lab managers Aivy David and Alejandro Ramirez-Apodaca. We thank the Xiaokun Shu lab and Shawn Douglas lab for lending us the equipment for directed evolution experiments. We thank Sarah Elmes for her help with FACS. FACS experiments were performed in the Laboratory for Cell Analysis, which is supported by a National Cancer Institute Cancer Center Support Grant (P30CA082103). S.Z. received support from the Chinese Scholar Council. This work is supported by grants from National Institutes of Health (R21GM129652, R01GM124334 and R01GM131641). B.H. is a Chan Zuckerberg Biohub Investigator.

## Supporting Information

**Fig. S1.**
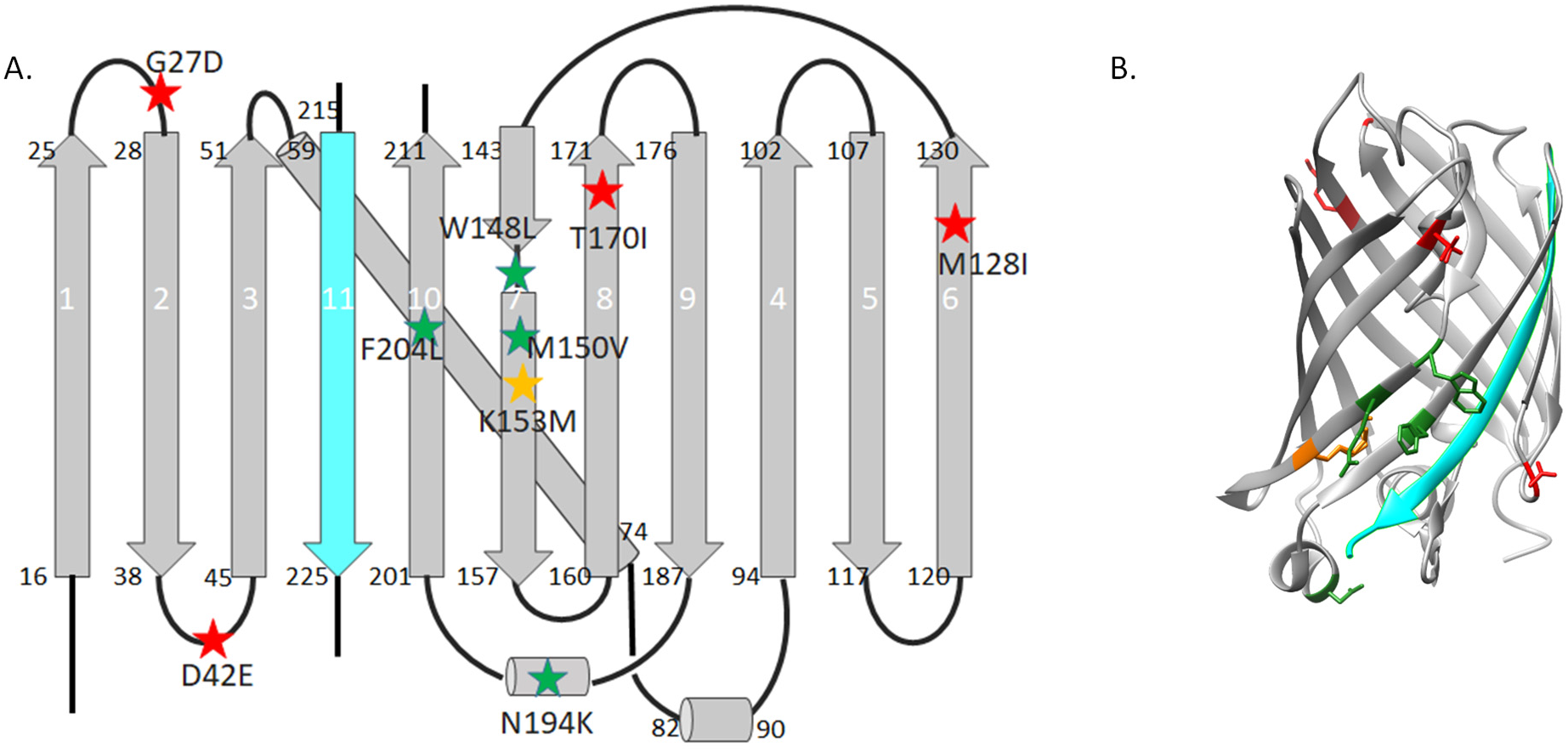
mNG mutations in 2D and 3D structure. A. 2D structure of mNG. β-sheets are shown as arrows and α-helices are shown as cylinders. Mutations are shown as stars. B. structure of mNG (PBD entry 5LTR). A-B. mNG3A colored in green, mNG3K in red, and shared mutation in orange, β-strand 11 labelled in cyan.

**Fig S2.**
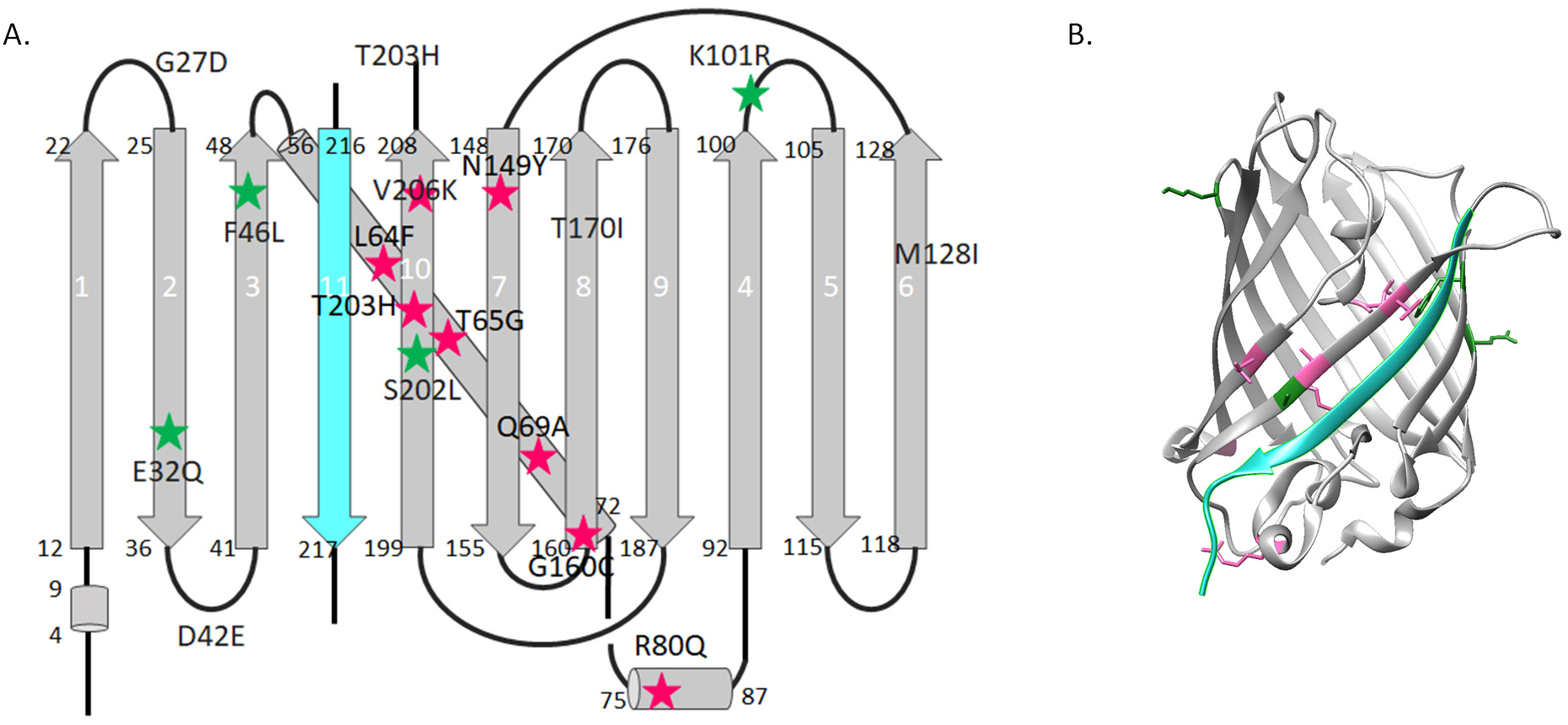
CloGFP mutations in 2D and 3D structure. A. 2D structure of sfGFP, β-sheets are shown as arrows and α-helices are shown as cylinders. Mutations are shown as stars. B, 3D structure of sfGFP from Protein Data Bank, ID 2B3P. A-B. mClover3 mutations in pink and CloGFP colored in green. β-strand 11 labelled in cyan.

**Fig S3.**
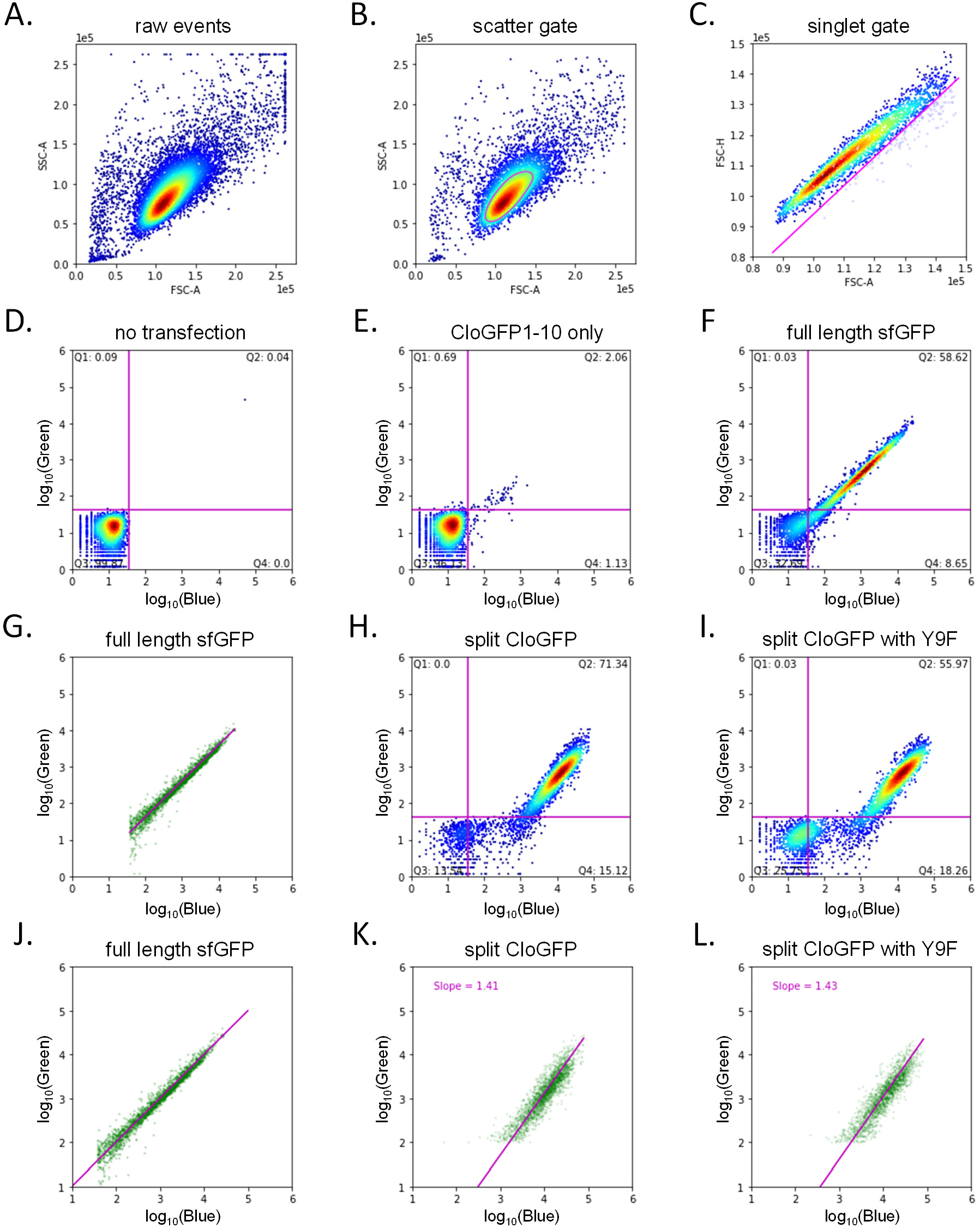
Flow Cytometry analysis of CloGFP_1-10_ background in the 405 nm excitation (blue) and 488 nm excitation (green) channels. A. Raw flow cytometry events. B. Scatter gating. C. Singlet gating. D. Negative control without transfection. E.Transfected with CloGFP_1-10_ only. F. Positive control with full length sfGFP. G. Fitting of full length GFP in BFP+ cells. H. CloGFP with wild type GFP_11_. I. CloGFP with Y9F mutant GFP_11_. J-L. Data from panels G-I rescaled for comparison.

**Fig S4.**
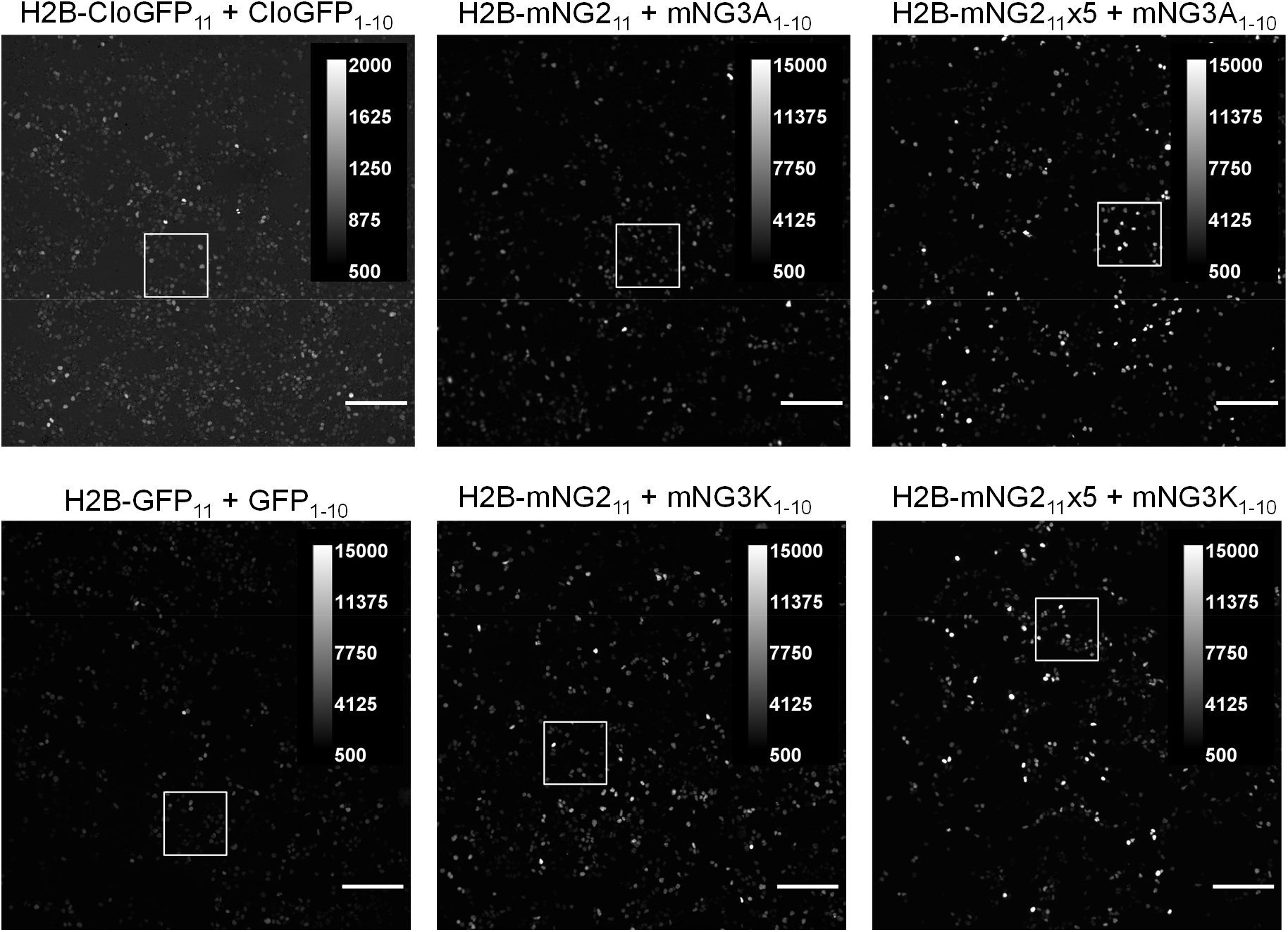
H2B labelling with new split proteins. Scale bars are 200 μm. Boxes indicate regions shown in Fig S5. Note that CloGFP is shown on a different intensity scale.

**Fig S5.**
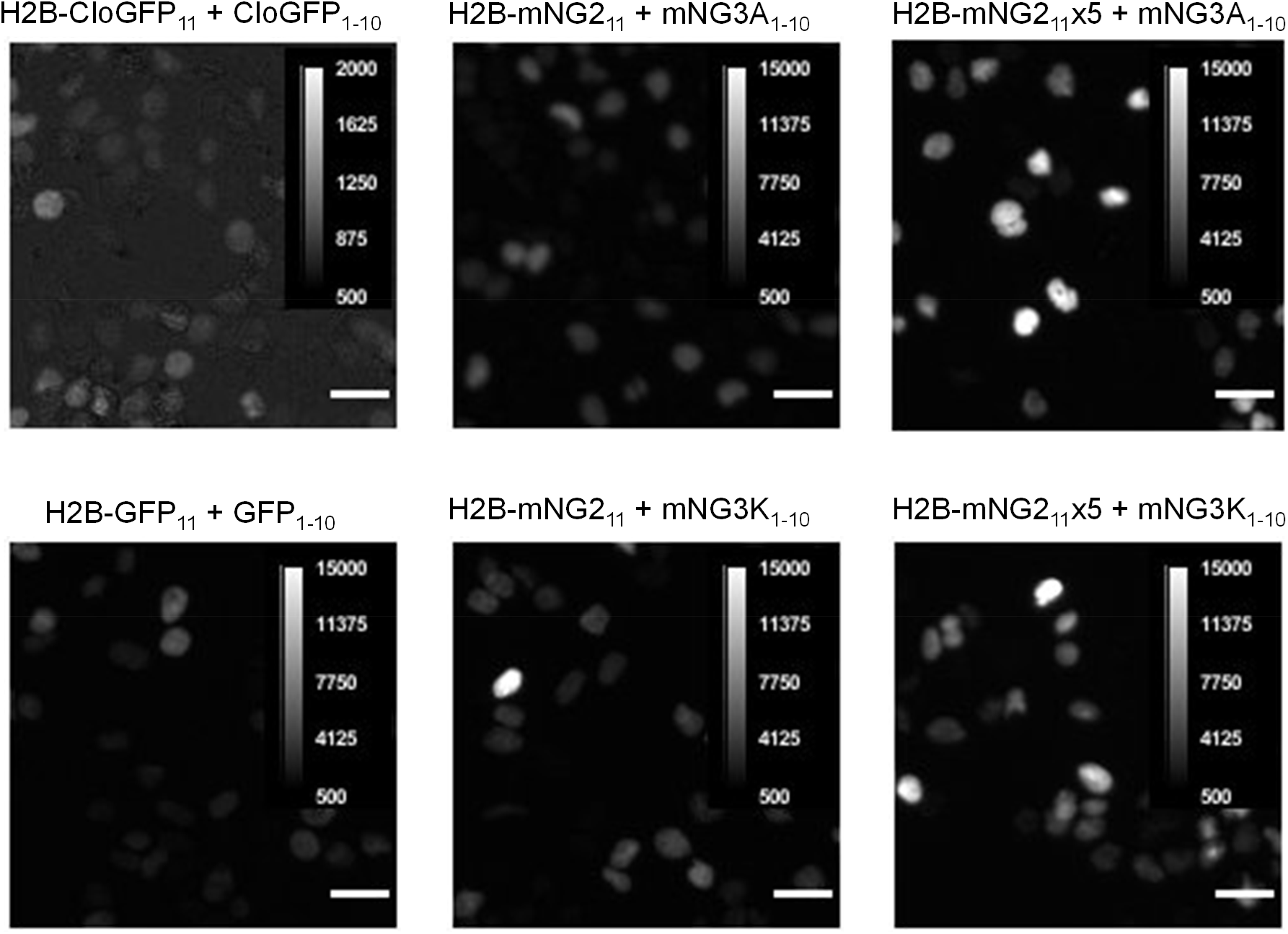
H2B labelling with new split proteins. Zoomed in images from the square box of Fig S4. White lines in the images are scale bars of 30 μm. Note the scale difference for the CloGFP compared with other images

**Table S1.** Mutation sites in each round of directed evolution for CloGFP.

| **Rounds** | **Genotype** | **Brightness** |
| --- | --- | --- |
| R0 | CloGFP0 | 805 |
| R1 | K101R | 3000 |
| R2 | K101R, S202L | 15300 |
| R3 | G32Q, F46L, K101R, S202L<br>Y9F(GFP11) | 34000 |
| R4 | G32Q, F46L, K101R, S202L<br>Y9F(GFP11) | 30000/30700/35000/36000 |

**Table S2.**
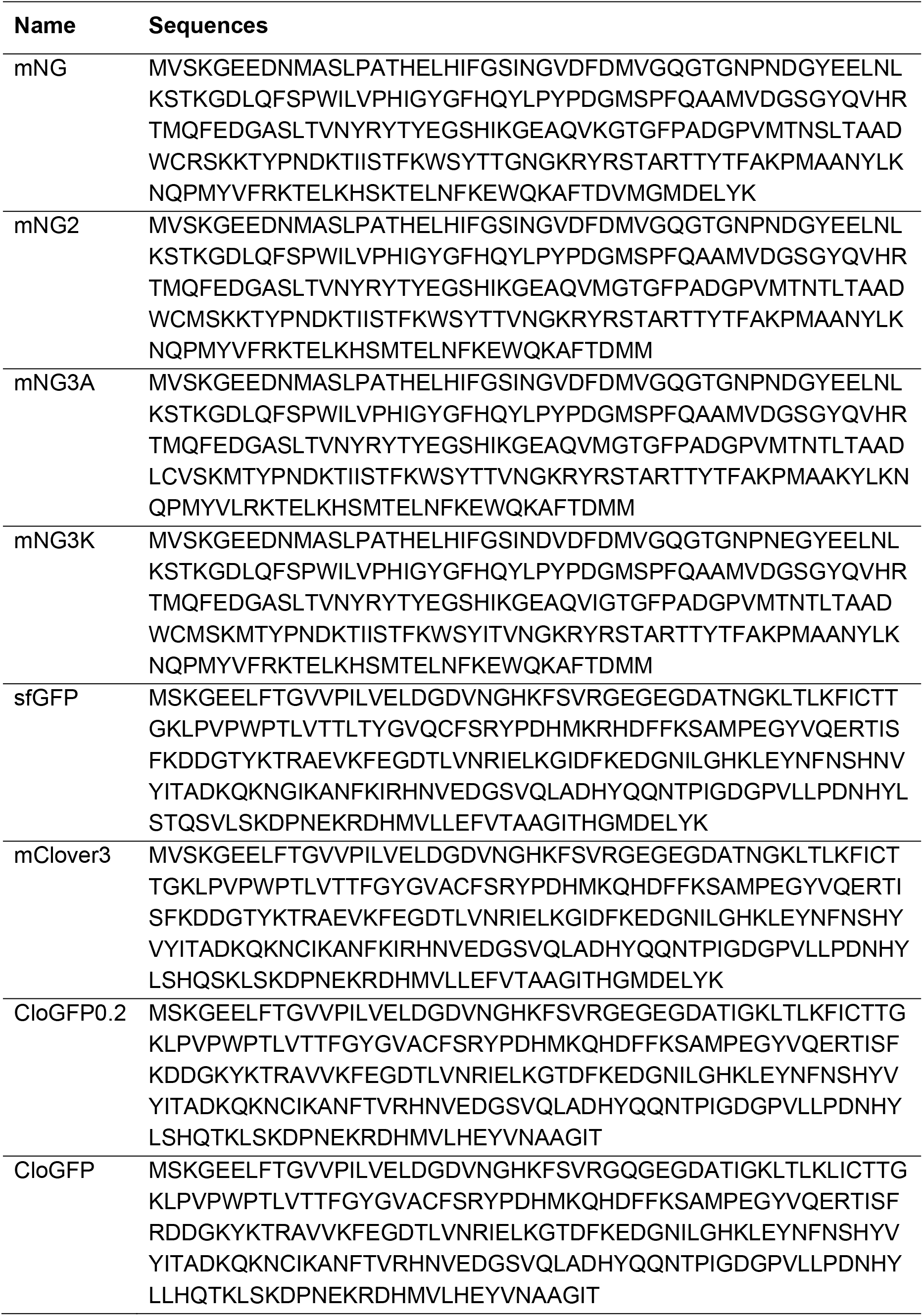
Protein sequences

## References

1. Cabantous S, Terwilliger TC, Waldo GS. Protein tagging and detection with engineered self-assembling fragments of green fluorescent protein. Nat Biotechnol. 2005 23(1):102–7.

2. Kamiyama D, Sekine S, Barsi-Rhyne B, Hu J, Chen B, Gilbert LA, et al. Versatile protein tagging in cells with split fluorescent protein. Nat Commun. 2016 18;7(1):11046.

3. Feng S, Sekine S, Pessino V, Li H, Leonetti MD, Huang B. Improved split fluorescent proteins for endogenous protein labeling. Nat Commun. 2017 8(1):370.

4. Feng S, Varshney A, Coto Villa D, Modavi C, Kohler J, Farah F, et al. Bright split red fluorescent proteins for the visualization of endogenous proteins and synapses. Commun Biol. 2019 2(1):1–12.

5. Kim J, Zhao T, Petralia RS, Yu Y, Peng H, Myers E, et al. mGRASP enables mapping mammalian synaptic connectivity with light microscopy. Nat Methods. 2011 9(1):96–102.

6. Batan D, Braselmann E, Minson M, Nguyen DMT, Cossart P, Palmer AE. A Multicolor Split-Fluorescent Protein Approach to Visualize Listeria Protein Secretion in Infection. Biophys J. 2018 115(2):251–62.

7. van der Schaar HM, Melia CE, van Bruggen JAC, Strating JRPM, van Geenen MED, Koster AJ, et al. Illuminating the Sites of Enterovirus Replication in Living Cells by Using a Split-GFP-Tagged Viral Protein. mSphere. 2016 1(4):e00104–16

8. Leonetti MD, Sekine S, Kamiyama D, Weissman JS, Huang B. A scalable strategy for high-throughput GFP tagging of endogenous human proteins. Proc Nat Acad Sci U S A. 2016 113(25):E3501–8.

9. Nagaraj N, Wisniewski JR, Geiger T, Cox J, Kircher M, Kelso J, et al. Deep proteome and transcriptome mapping of a human cancer cell line. Mol Sys Biol. 2011 7:548.

10. Shaner NC, Lambert GG, Chammas A, Ni Y, Cranfill PJ, Baird MA, et al. A bright monomeric green fluorescent protein derived from Branchiostoma lanceolatum. Nat Methods. 2013 10(5):407–9.

11. Ah K, A B, Mp F, Sc R, Inak A, M H. Evolving Accelerated Amidation by SpyTag/SpyCatcher to Analyze Membrane Dynamics. Angew Chem Int Ed. 2017 56(52):16521–16525

12. Lam AJ, St-Pierre F, Gong Y, Marshall JD, Cranfill PJ, Baird MA, et al. Improving FRET dynamic range with bright green and red fluorescent proteins. Nat Methods. 2012 9(10):1005–12.

13. Bajar BT, Wang ES, Lam AJ, Kim BB, Jacobs CL, Howe ES, et al. Improving brightness and photostability of green and red fluorescent proteins for live cell imaging and FRET reporting. Sci Rep. 2016 6(1):20889.

14. Kerppola TK. Visualization of molecular interactions by fluorescence complementation Nat. Rev. Mol. Cell Biol. 2006 7:449–456.

15. English JG, Olsen RHJ, Lansu K, Patel M, White K, Cockrell AS, et al. VEGAS as a Platform for Facile Directed Evolution in Mammalian Cells. Cell. 2019 178(3):748–761.e17.

16. Erdogan M, Fabritius A, Basquin J, Griesbeck O. Targeted In Situ Protein Diversification and Intra-organelle Validation in Mammalian Cells. Cell Chem Biol. 2020 27(5):610–621.e5.

17. Chen H, Liu S, Padula S, Lesman D, Griswold K, Lin A, et al. Efficient, continuous mutagenesis in human cells using a pseudo-random DNA editor. Nat Biotechnol. 2020 38(2):165–8.

